# Genomic Analysis of a Transcriptional Networks Directing Progression of Cell States During MGE development

**DOI:** 10.1101/329797

**Authors:** Magnus Sandberg, Leila Taher, Jiaxin Hu, Brian L. Black, Alex Nord, John L.R. Rubenstein

## Abstract

**Background:** Homeodomain (HD) transcription factor (TF) NKX2-1 critical for the regional specification of the medial ganglionic eminence (MGE) as well as promoting the GABAergic and cholinergic neuron fates via the induction of TFs such as LHX6 and LHX8. NKX2-1 defines MGE regional identity in large part through transcriptional repression, while specification and maturation of GABAergic and cholinergic fates is mediated in part by transcriptional activation via TFs such as LHX6 and LHX8. Here we analyze the signaling and TF pathways, downstream of NKX2-1, required for GABAergic and cholinergic neuron fate maturation.

**Methods:** Differential ChIP-seq analysis was used to identify regulatory elements (REs) where chromatin state was sensitive to change in the *Nkx2-1*cKO MGE at embryonic day (E) 13.5. TF motifs in the REs were identified using RSAT. CRISPR-mediated genome editing was used to generate enhancer knockouts. Differential gene expression in these knockouts was analyzed through RT-qPCR and in situ hybridization. Functional analysis of motifs within hs623 was analyzed via site directed mutagenesis and reporter assays in primary MGE cultures.

**Results:** We identified 4782 activating REs (aREs) and 6391 repressing REs (rREs) in the *Nkx2-1* conditional knockout (*Nkx2-1*cKO) MGE. aREs are associated with basic-Helix-Loop-Helix (bHLH) TFs. Deletion of hs623, an intragenic *Tcf12* aRE, caused a reduction of *Tcf12* expression in the sub-ventricular zone (SVZ) and mantle zone (MZ) of the MGE. Mutation of LHX, SOX and octamers, within hs623, caused a reduction of hs623 activity in MGE primary cultures.

**Conclusions:** *Tcf12* expression in the sub-ventricular zone (SVZ) of the MGE is mediated through aRE hs623. The activity of hs623 is dependent on LHX6, SOX and octamers. Thus, maintaining the expression of *Tcf12* in the SVZ involves on TF pathways parallel and genetically downstream of NKX2-1.

## BACKGROUND

Transcription factors (TFs) direct cell fate determination and differentiation through binding to a genomic network consisting of regulatory elements (REs) such as promoters and enhancers. By analyzing epigenetic modifications and transcriptional changes in TF knockouts, we have started to uncover the genomic networks and molecular mechanisms that direct brain development [1]. In-depth understanding of the genetically encoded wiring of the brain is important as perturbation of transcription pathways is implicated in disorders such as autism and intellectual disability [2]. Distantly acting REs have been identified based on conservation and activity [3,4]. Their spatial activity and dynamic genomic contacts can be predicted using a combination of TF binding profiling, genome-wide 3D chromosome organization mapping and CRISPR/Cas9 editing [5–10]

Mouse genetic experiments have elucidated the functions of many TFs in the development of the subpallial telencephalon [11,12]. These studies show that the HD protein NKX2-1 is required for regional specification of the MGE by repressing alternative identities, as well as promoting GABAergic and cholinergic cell fates via the induction of TFs such as LHX6 and LHX8 [13–17]. By integrating genomic data with mouse genetics, we confirmed the repressive function of NKX2-1, however its role in transcriptional activation remains unclear. Moreover, additional data suggests that genes genetically downstream of NKX2-1, such as LHX6 and LHX8, are responsible for the loss of gene expression observed in the *Nkx2-1*cKO [18,19]. Altogether, the genetic program and molecular mechanisms responsible for promoting GABAergic and cholinergic neuron phenotypes, downstream of NKX2-1 remains largely unexplored.

To investigate the signaling pathways of MGE development downstream of NKX2-1, we extended our earlier analysis of the genomic network directing MGE development that is altered in the *Nkx2-1* mutant. First we evaluated all loci that showed an epigenetic change, independent of NKX2-1 binding. Via an epigenomic analysis of the NKX2-1 mutant MGE we characterized a large set REs that are implicated in mediating transcriptional repression and activation. Using a combination of genomics, *de novo motif* analysis, CRISPR engineering and primary culture assays we characterize REs and TFs central to patterning of the subpallial telencephalon and promoting MGE characteristics. Gene ontology (GO) analysis showed an enriched association of REs activating transcription (aREs) with E-box binding basic-Helix-Loop-Helix (bHLH) TFs. Using CRISPR engineering we deleted hs623, an intronic aRE of the *Tcf12* gene which encodes a bHLH TF. Deletion of hs623 reduced *Tcf12* expression in the MGE. *De novo* motif analysis combined with TF motif mutations, showed that OCT/POU and SOX motifs are required for hs623’s ability to promote transcription in the MGE.

## METHODS

### Mice

The *Nkx2-1*cKO was earlier described in Sandberg et al. 2016 [18] and generated using mice strains previously reported: *Nkx2-1f/f* [20], *Olig2-tva*-Cre [21] and *AI14* Cre-reporter [22]. All experiments with animals complied with federal and institutional guidelines and were reviewed and approved by the UCSF Institutional Animal Care and Use Committee.

### Generation of *hs623* deletion

The *hs623*^*Tm1*^ allele was generated by CRISPR-mediated genome editing, using established methods [23]. Guide RNAs sgRNA-hs623-1, 5′-GTTTAGTTTTGCTCATACCA(TGG)-3′ and sgRNA-hs623-2, 5′-ATGGTTTCTGTGATCGTAAT(TGG)-3′ (protospacer-adjacent motif [PAM] sequence indicated in parentheses) were transcribed *in vitro* using the MEGAshortscript T7 kit (Life Technologies, AM1354) and subsequently purified using the MEGAclear kit (Life Technologies, AM1908). The two guide RNAs were designed to delete a 737bp intronic region within *Tcf12* [mm9; chr9:71822812-71823548]. The purified sgRNAs were co-injected into the cytoplasm of fertilized mouse oocytes with *in vitro* transcribed Cas9 mRNA using standard transgenic procedures as previously described [24]. F0 transgenic founders were identified by PCR screening using hs623-KO-F, 5’-GTGGCTGATGATGTGCTCTGA-3’ and hs623-WT-KO-R, 5’-CTCCATCAGGTTCTTGCCCC-3’ to identify the *hs623* null alleles (KO = 250bp, WT = 1008bp) and hs623-WT-F, 5’-GTGGCTGATGATGTGCTCTGA-3’ and hs623-WT-KO-R to identify the *hs623*-WT allele (462bp). Four independent F0 founders were each outcrossed to wild type mice, and F1 offspring were used for subsequent *hs623*^*Tm1*^ intercrosses to generate *hs623*-null mice.

### Histology

Immunofluorescence was performed on 16 µm cryosection as previously described [25]. *In situ* hybridization was performed as previously described [26]. The following primers were used generate the templates used for the *in situ* probes: Tcf12_F, TCTCGAATGGAAGACCGC; Tcf12_R, CTCCCTCCTGCCAGGTTT

### Dissection of Embryos

RT-qPCR and primary culture experiments were performed on E13.5 micro-dissected MGE. All MGE dissections were performed as follows; the dorsal boundary was defined by the sulcus separating lateral ganglionic eminence (LGE) and MGE. The caudal end of the sulcus defined the caudal boundary. Septum was removed.

### Gene expression analysis in *hs623*KO

To assay differential gene expression in the *hs623*KO RNA was purified using RNEasy Mini (Qiagen) and cDNA was generated using Superscript III^®^ First-Strand Synthesis System for RT-PCR (Invitrogen). RT-qPCR analysis was performed on a 7900HT Fast Real-Time PCR System (Applied Biosystems) using SYBR GreenER qPCR SuperMix (Invitrogen, Cat. No. 11760-100). Unpaired t-test was used to test significance in gene expression between *hs623*WT and *hs623*KO using SDHA as internal control [27,28].

Sequences of RT-qPCR primers used:

SDHA-F, GCTCCTGCCTCTGTGGTTGA

SDHA-R, GCAACACCGATGAGCCTG

Mns1_ctrl_1F, CTGCTGCTCCGGAAGACG

Mns1_ctrl_1R, TTTTGGTCGCCATCTCGGTT

Myzap_ctrl_2F, TCGAAAGGAAAGATCAGCCTCC

Myzap_ctrl_2R, TCTGATCTTCGCACCACACC

Zfp280d_ctrl_1F, CCCCAGCTCTCATTCAAGAGG

Zfp280d_ctrl_1R, TTCAGGCAGCGTTGACTTGT

TCF12_v1/2-F2, GCTTGTCCCCAACACCTTTC

TCF12_v1/2-R2, TGACAGCCTGAGAGTCCAGA

TCF12_v1/3-F4, TACCAGTCAGTGGCCCAGAG

TCF12_v1/3-R4, AATGCTCGTGAAGTTGCTGC

TCF12_v1/3-F5, TCCCTGGAATGGGCAACAAT

TCF12_v1/3-R5, TCACGGTTGAAATCGTCAGA

### Site-directed mutagenesis of TF binding motifs in *hs623*

To study the requirement TF motifs for hs623 activity LHX6, SOX and octamers were mutated in pCR-Blunt II-TOPO, sequence verified and sub-cloned into a pGL4.23-Luciferase reporter with a minimal β-globin-promoter using BglII and XhoI [18]. Following primers were used to generate the different hs623 luciferase reporters:

hs623-mut-site#1-R, cgttgctgacaaggctgttttttacagaaattgatgctgagttc

hs623-mut-site#1-F, agccttgtcagcaacgtgattattcaaac

hs623-mut-site#2-F, gatgtgctctgatatgaaaaaagtcattaggtagaatgaatag

hs623-mut-site#3-F, gatgtgctctgatatgtaattagaaaaaaggtagaatgaatag

hs623-mut-site#2 and 3-F, gatgtgctctgatatgaaaaaagaaaaaaggtagaatgaatag

hs623-mut-site#2 and/or 3-R, atatcagagcacatcatcagccacattc

hs623-mut-site#4-F, gattattcaaacaactcttttttttgttaatgagg

hs623-mut-site#4-R, gagttgtttgaataatcacgttgctgac

hs623-mut-site#5-F, ctcatgcaaatgaaaaagaggccttatttgc

hs623-mut-site#5-R, atttgcatgagttgtttgaataatc

hs623-mut-site#4 and 5-F, caaacaactcttttttttgaaaaagaggccttatttgc

hs623-mut-site#4 and 5-R, use “

hs623-mut-site#4-R” for PCR.

hs623-mut-site#6-F, gttaatgaggccttaaaaaaatatttattttttcc

hs623-mut-site#6-R, ggcctcattaacatttgcatgagttgtttg

hs623-mut-site#4 and 6-F, caactcttttttttgttaatgaggccttaaaaaaatatttattttttcc

hs623-mut-site#4 and 6-R, cattaacaaaaaaaagagttgtttgaataatcac

hs623-mut-site#4,5 and 6-F, caactcttttttttgaaaaagaggccttaaaaaaatatttattttttcc

hs623-mut-site#4, 5 and 6-R, ggcctctttttcaaaaaaaagagttgtttgaataatc

hs623-mut-site#7-F, gcaacgtgattattcccccccctcatgcaaatg

hs623-mut-site#7-R, gaataatcacgttgctgacaagg

hs623-mut-site#4 and 7-F, gtgattattcccccccctcttttttttgttaatgagg

hs623-mut-site#4 and 7-R, use hs623-mut-site#7-R

### Analysis of hs623 activity in MGE primary MGE cultures

MGE tissue was dissected from E13.5 embryos, triturated and plated onto 24-well plates (1 embryo / 2wells). Primary cultures were transfected with a total of 500ng DNA using Lipofectamin 2000 (Thermo Fisher) and cultured in Neurobasal Medium (Thermo Fisher) supplemented with 0.5% Glucose, GlutaMAX (Thermo Fisher Scientific) and B27 (Thermo Fisher Scientific). Luciferase assays were performed 48h after transfection using Dual Luciferase Reporter Assay System (Promega). Unpaired t-test was used to test significance between the variants of hs623.

### ChIP-Seq Computational Analysis

Differential ChIP-seq analysis was performed as described in Sandberg et al. 2016 [18]. After differential H3K4me1, H3K27ac and H3K27me3 analysis we merged overlapping sites. Only merged sites that were enriched in H3K4me1 relative to the input datasets and for which the difference in enrichment between *Nkx2-1* WT and cKO was not significant (for at least one of the sites among the merged sites) were further considered. Of those, merged sites overlapping with blacklisted genomic regions (http://mitra.stanford.edu/kundaje/akundaje/release/blacklists/mm9-mouse/mm9-blacklist.bed.gz) and RepeatMasker annotation (http://hgdownload.cse.ucsc.edu/goldenPath/mm9/database/chr*_rmsk.txt.gz) as well as those exceeding 5000bp were excluded. We defined aREs based on the following two criteria; 1) more H3K27ac (WT) and no increase in H3K27me3 (WT), H3K27ac (*Nkx2-1*cKO) and H3K4me1 (*Nkx2-1*cKO), 2) more H3K27me3 (*Nkx2-1*cKO) and no increase in H3K27ac (*Nkx2-1*cKO), H3K4me1 (*Nkx2-1*cKO) and H3K27me3 (WT). We defined rREs based on the following two criteria; 1) a RE with more H3K27ac (*Nkx2-1cKO)* and no increase in H3K27me3 (*Nkx2-1*cKO), H3K27ac (WT) and H3K4me1 (WT), 2) more H3K27me3 (*WT)* and no increase in H3K27ac (WT), H3K4me1 (*WT)* and H3K27me3 (*Nkx2-1*cKO).

### *De novo* motif analysis

Motif analysis performed using RSAT [29] with default settings and genomic, aREs or rREs as background.

## RESULTS

### Identification of the genomic regulatory network directing MGE identity

We have previously shown that the combined binding of NKX2-1 and LHX6 is a predictive indicator of REs that mediate transcriptional activation in the subventricular (SVZ) and mantle zone (MZ) of the MGE in the developing subpallial telencephalon [18]. There is evidence that NKX2-1 generally acts as a repressor in MGE progenitors (in the ventricular zone [VZ]), whereas LHX6, and potentially other TFs and signaling pathways, some of which are genetically downstream of NKX2-1, are important for activating transcription in the SVZ and MZ of the MGE [18,30]. By studying aREs, we aimed to further explore the molecular mechanisms underlying the transcriptional network directing differentiation of the secondary progenitors in the SVZ.

First we identified aREs and rREs by assessing the genome-wide changes of the two histone marks H3K27ac and H3K27me3 at H3K4me1 positive REs comparing the WT and *Nkx2-1*cKO MGE [18]. We defined aREs based on the following two criteria; 1) reduced H3K27ac and, 2) increased H3K27me3 in the *Nkx2-1*cKO. We defined rREs based on the following two criteria; 1) increased H3K27ac and, 2) reduced H3K27me3 in the *Nkx2-1*cKO (see Methods).

Based on these criteria we identified 4782 aREs and 6391 rREs in the *Nkx2-1*cKO. See Additional file 1 for a complete list of aREs and rREs. To analyze the *in vivo* activity patterns of the aREs and rREs we examined transgenic enhancer activity patterns of E11.5 forebrain enhancer activity patterns available in the VISTA database (see VISTA data base; https://enhancer.lbl.gov/) [7]. The activities of rREs were highest in cortex (62%) and LGE and dorsal MGE (52%) and lowest in the ventral MGE (24%)(Figure 1A and 1B [hs848, hs1172 and hs1187]). Note, hs1187 is active in the, NKX2-1 negative, dorsal most part of the MGE illustrating the repressing activity of NKX2-1 on this type of rREs [31]. In contrast, aREs have the highest activities in the MGE (53%) when compared to their activities in the LGE (50%) and cortex (41%) (Figure 1A and 1B [hs676, hs957 and hs1041]). We also found a higher activity of MGE positive aREs in the SVZ and MZ compared to the VZ, consistent with our previous results for NKX2-1 bound aREs and rREs (Figure 1B and 1C) [18]. See Additional file 1 for a full list of aREs and rREs VISTA transgenics.

**Figure 1.**
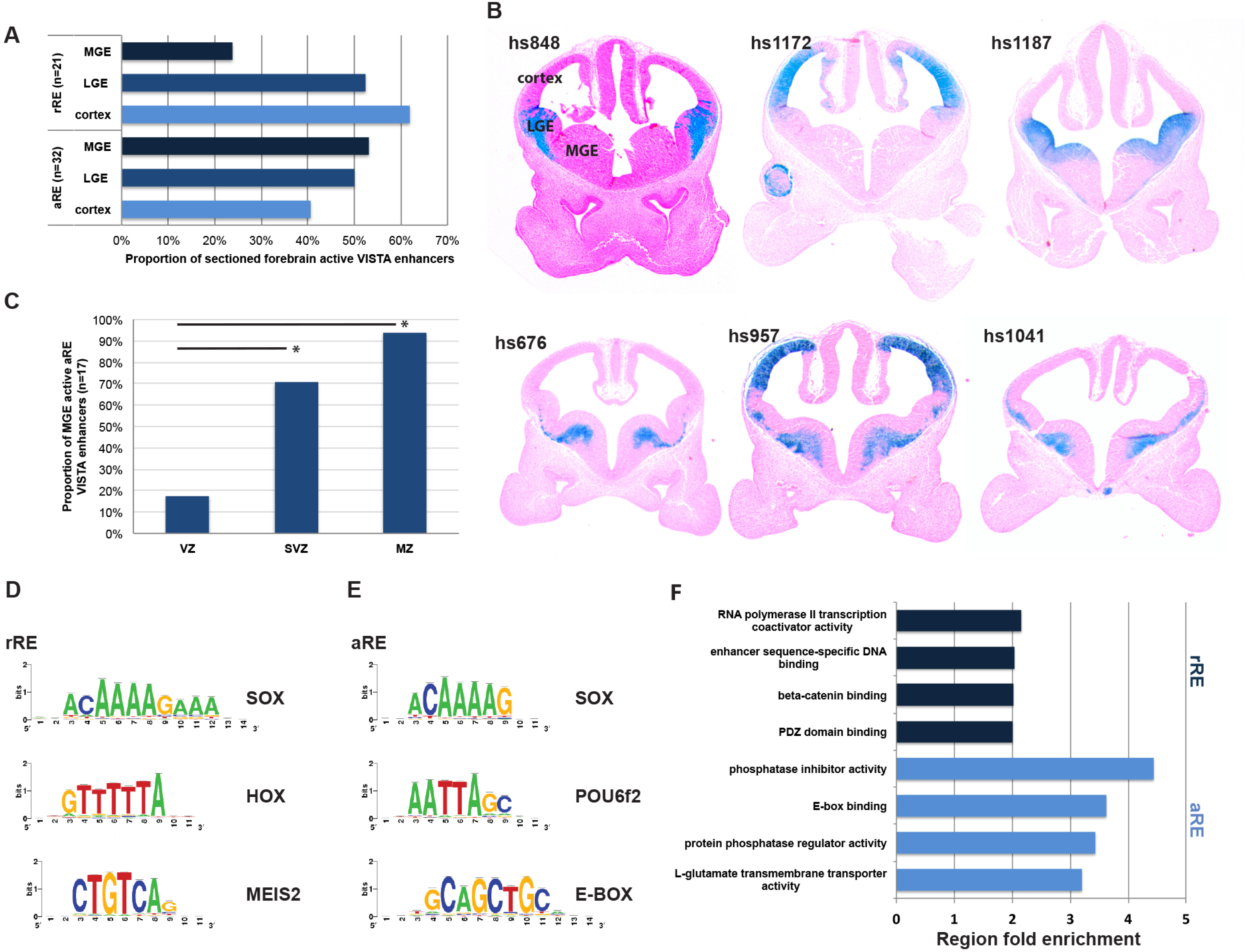
Characterization of aREs and rREs in E13.5 MGE. (A) Proportion of aREs and rREs active in MGE, LGE and cortex. (B) Sections of transgenic embryos (from the VISTA browser) showing *in vivo* activity of rREs (hs848, hs1172 and hs1187) and aREs (hs676, hs957 and hs1041) at E11.5. (C) VZ, SVZ, and MZ activity of aREs in the MGE at E11.5. Chi-square test was used to test significance between the groups: *p < 0.05. (D-E) Manually curated list of *de novo* motifs and potential TF recognizing the motifs in rREs and aREs. (F) Enriched gene ontology annotations of putative aRE target genes.

To identify TFs motifs enriched in the aRE and rREs we performed a *de novo* motif discovery using RSAT [29]. This analysis showed a number of motifs enriched in both aREs and rREs such as SOX motifs, homedomain binding motifs (HOX and POU6f2) and motifs recognized by zinc finger TFs (e.g. SP1 and ZNF384) (Figure 1D and 1E). Additional analysis identifying motifs differentially enriched between aREs and rREs showed that aREs have a high frequency of E-boxes (Figure 1E). Interestingly, we found that rREs are enriched in motifs consistent with the binding site of the TF MEIS2 (Figure 1D). The *Meis2* gene is repressed by NKX2-1, and in turn, its RNA is strongly up-regulated in the MGE of the *Nkx2-1*cKO [18]. These data suggest that *Meis2* is central to activating a genomic network promoting LGE and caudal ganglionic eminence (CGE) characters (through rREs) in the *Nkx2-1*cKO MGE.

We then examined enrichment of annotation terms among the aREs and rREs candidate target genes using GREAT [32]. Top-ranked GO terms for rREs target genes were associated with WNT signaling (beta-catenin binding and PDZ domain binding), transcriptional regulation (such as RNA polymerase II transcription co-activator activity), and enhancer sequence-specific DNA binding (Figure 1F). Looking specifically at the associated genes for the rREs containing MEIS2 binding motifs we found several genes (*Isl1, Ebf1, Tle4, Zfp503, Efnb1* and *Efnb2*) with higher expression in the LGE and CGE than the MGE. These findings support the hypothesis that MEIS2 directs LGE and CGE identities. The top-ranked GO terms for aREs target genes were associated with phosphatase activity, E-box binding proteins, L-glutamate transmembrane transporter activity and transmembrane-ephrin receptor activity [32](Figure 1F). Two E-box binding TFs, *Tcf4* and *Tcf12*, which are in the region of a large number of aREs, have reduced MGE SVZ and MZ expression in the *Nkx2-1*cKO [18]. In combination with the high frequency of E-boxes in aREs, our data suggests that *Tcf4* and *Tcf12* are components of the genomic network regulating gene expression in secondary progenitors of the MGE that are genetically downstream of NKX2-1.

### *In vivo* characterization of *hs623* in the MGE of the forebrain

To learn more about the *Tcf12* expression and the transcriptional pathways integrated in the aRE network downstream of NKX2-1, we examined aRE *hs623*, a highly evolutionarily conserved 914 base pair (bp) sequence that is in an intron of the *Tcf12* locus (Figure 2A and 2B). A previous transgenic study show that *hs623* drives *LacZ* expression at E11.5 [33]. The *hs623* transgene is active in the forebrain, hindbrain and the spinal cord (Figure 2C-2E). A coronal section through the telencephalon shows that *hs623* activity is restricted to the SVZ and MZ of the MGE, and perhaps labels cell tangentially migrating into the LGE (Figure 2E). This pattern of activity is supported by histone ChIP-seq analysis of the MGE showing that this locus has histone modifications that are characteristic of active enhancer elements (Figure 2A [H3K4me1 + and H3K27ac +] and 2B). Of note, ChIP-seq analysis of the MGE *Nkx2-1*cKO shows reduced H3K27ac, providing evidence that the activity of the locus is dependent on the activity of the NKX2-1 and / or its target TFs, as reported earlier (Figure 2B) [18].

**Figure 2.**
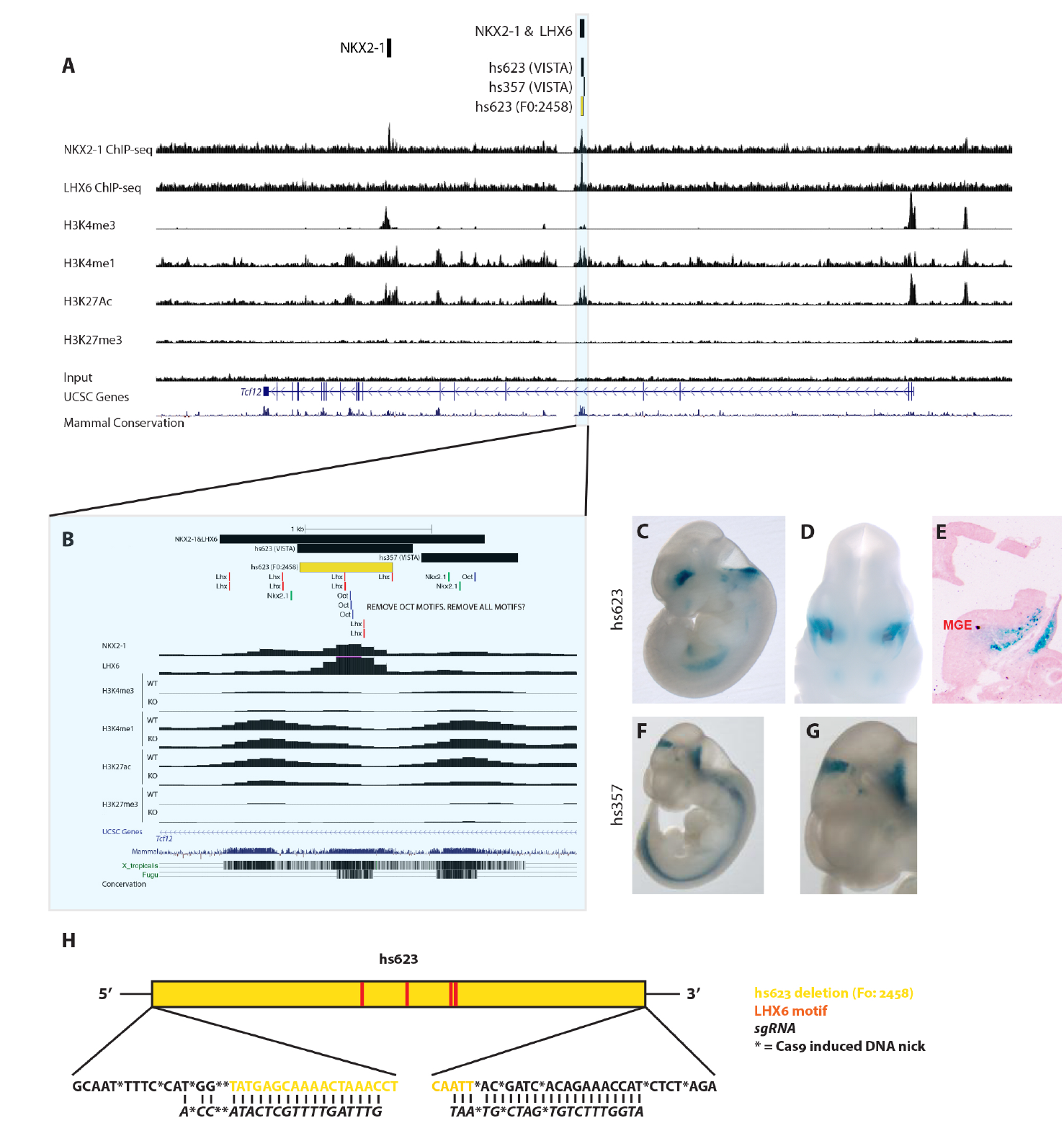
Deletion of *cis*-regulatory element hs623 *in vivo*. (A) Genomic region of the *Tcf12* locus with the ChIP-seq datasets and genomic features shown; NKX2-1 ChIP-seq, LHX6 ChIP-seq, H3K4me3, H3K4me1, H3K27ac, H3K27me3, UCSC genes and mammalian conservation. Histone 3 modifications in MGE at E13.5. Hs623 region framed and highlighted in blue. (B) Higher resolution view of the hs623 region with the same ChIP-seq datasets as in Figure 2A. Called NKX2-1 & LHX6 binding region, VISTA regions, deleted hs623 region (yellow) and NKX2-1, LHX6 and OCT consensus motifs labeled at the top of the browser. (C-G) VISTA database transgenic embryos showing *in vivo* activity of *hs623* and *hs357* at E11.5. (H) Schematic description of the generated hs623 deletions (5 founders). The distribution of LHX6 consensus motifs in hs623 are indicated. Founder 2458 was used for the analysis presented in this paper.

### Motif logic direct region specific transcriptional activity

*Hs623* is flanked by two highly conserved regions and the activity of one of the regions (*hs357)* has been tested *in vivo* [33]. Similar to *hs623, hs357* is active in the spinal cord, but unlike *hs623* it is active in the pretectum and it lacks activity in the telencephalon, including the MGE (Figure 2F and 2G). Therefore, despite the close proximity of *hs623* and *hs357*, they show differences in regional activity, suggesting that their regional activities are more likely due to differences in their primary nucleic acid sequence rather then their genomic location. Consistent with the lack of MGE activity, *hs357* lacks LHX6 consensus motifs (Figure 2B). On the other hand, *hs623* has four LHX6 consensus motifs (Figure 2H). Surprisingly, even though *hs623* has NKX2-1 binding, it contains no NKX2-1 consensus motifs. However, when extending the sequence analysis to the regions flanking *hs623,* we find three NKX2-1 consensus motifs, two within *hs357* (Figure 2B). This could explain why we detect NKX2-1 binding a wide region that covers both *hs623* and *hs357*.

### CRISPR / Cas9 mediated deletion of *hs623 in vivo*

To functionally test the requirement of *hs623 in vivo*, we deleted *hs623* using CRISPR / Cas9 (see VISTA database; http://enhancer.lbl.gov). A pair of sgRNAs were designed to delete the 734bp core sequence of *hs623*, which has NKX2-1 and LHX6 binding (Figure 2B and 2H). Micorinjection of sgRNAs and Cas9 generated a total of 22 pups. 23% (5 of 22) of the pups carried the desired *hs623* deletions and the induced DNA breaks were distributed within 20bp of the predicted cutting site (5’ and 3’ of hs623, Figure 2H). To minimize potential off target effects we outcrossed the F0 transgenic founders to wild-type CD1 mice. Four of five founders were fertile and generated a F1 generation; these animals were intercrossed to generate homozygous F2 *hs623KO* animals. *Hs623KOs* in the F2 generation were produced at Mendelian Ratios showing that the enhancer deletion was viable (10 [WT]:19 [HET]:7 [KO], n = 3 litters; 𝒳^2^ = 0.611; df = 2; *p* = 0.7367). Due to the overall similarity of the four fertile founders we decided to focus the following analysis on one of the founders (F0: 2458, Figure 2H).

### Deletion of *hs623* reduces *Tcf12* mRNA levels in the SVZ of MGE

*Hs623* is a *Tcf12* intragenic RE that in transgenic assays activates transcription in the SVZ of the MGE (Figure 2C-E). As noted above, its activity is partly dependent on NKX2-1 activity and *Tcf12* transcription is specifically reduced in the SVZ of the MGE in the *Nkx2-1*cKO (Figure 2B) [18]. Together, these data suggest that *hs623* could be a RE activating *Tcf12* transcription in the MGE. To test this hypothesis, we performed RTqPCR on the MGE from *hs623WTs* and *hs623KOs* at E13.5. Primers were designed to target all known mouse protein-coding and non-protein-coding genes in the NCBI RNA reference sequences collection that are found 450 kb up-and downstream of *hs623* (Figure 3A). From the RTqPCR we found no significant difference in the expression of the following genes in this region: *Myzap, Cgln1, Zfp280d* and *Mns1* (Figure 3B). *Tcf12* RNAs include a variety of splice variants. Because of this we designed three separate primer pairs to specifically interrogate the different splice variants of *Tcf12* (Figure 3A). We found a reduction in the expression of the short isoforms of *Tcf12* isoform 3 and 4 (Figure 3B, see *Tcf12*_v1/3-4 and *Tcf12*_v1/3-5). Notably, we did not find any difference in the expression levels of the longer isoforms 1 and 2 of *Tcf12* (Figure 3B, see *Tcf12*_v1/2-2). Together, these results show that *Tcf12* transcription in the MGE is enhanced by *hs623*.

**Figure 3.**
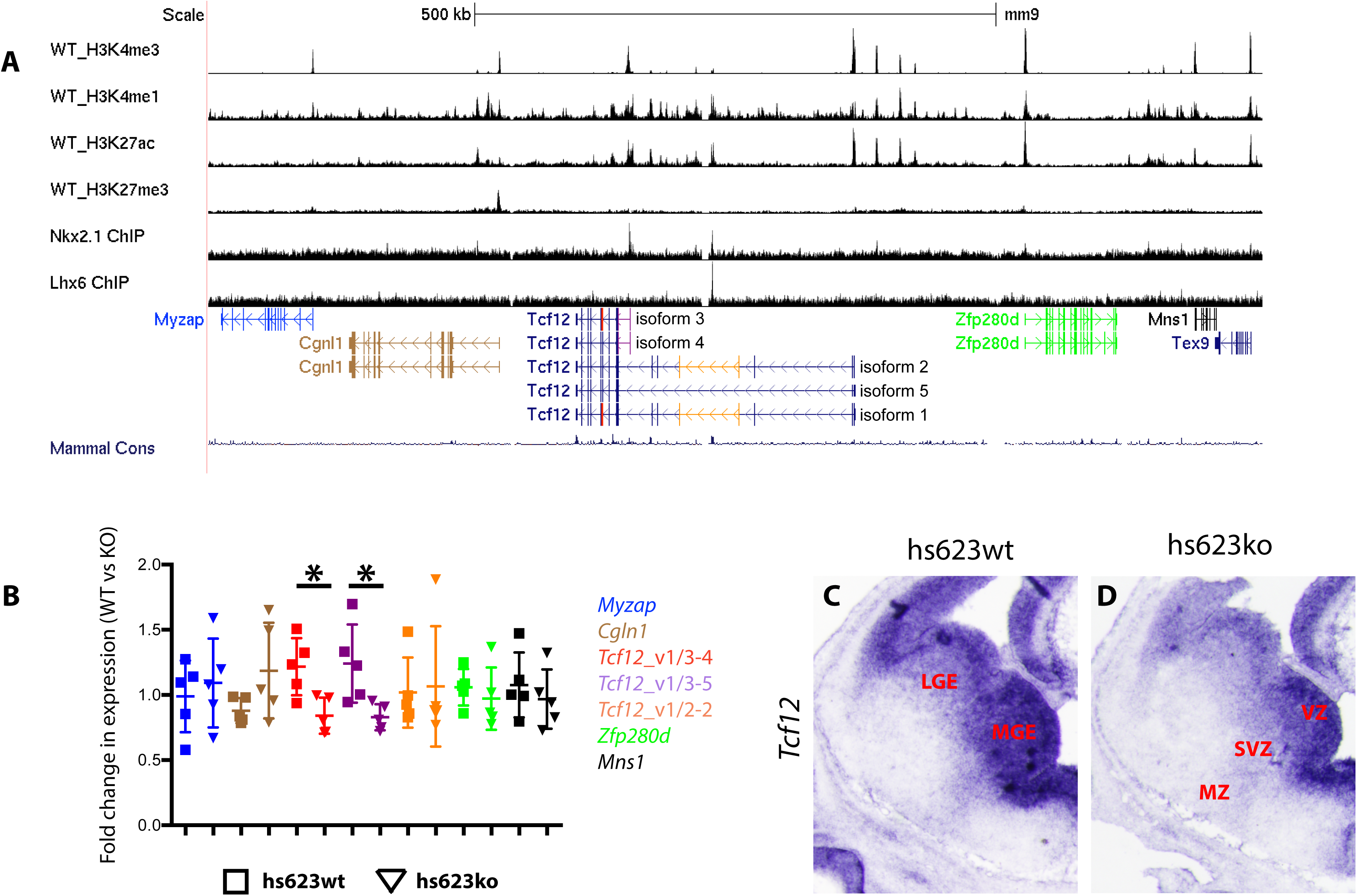
Reduced *Tcf12* expression in the *hs623*KO. (A) *Tcf12* locus with neighboring genes. (B) qPCR analysis of *Mns1, Myzap, Cgln1, Zfp280d, Tcf12* isoforms 1 and 3 (*Tcf12*_v1/3-4), *Tcf12* isoforms 3 and 4 (*Tcf12*_v1/3-5), *Tcf12* variant 1 and 2 (*Tcf12*_v1/2-2) on WT and *hs623*KO MGE at E13.5. The colors used in the table correlate to the specific target regions indicated in Figure 3A. (C-D) *In situ* hybridization analysis of *Tcf12* in WT (C) and *hs623*KO (D) basal ganglia at E13.5. Unpaired t-test was used to test significance between the groups: *p < 0.05.

To obtain spatial information about the reduction of *Tcf12* within the MGE we compared the distribution of *Tcf12* RNA between WT and *hs623KO* telencephalon at E13.5 using *in situ* RNA hybridization. Normally, *Tcf12* is broadly expressed in the VZ in the pallium and subpallium. In the ganglionic eminences *Tcf12* is also expressed in the SVZ and MZ, with a markedly higher expression in the MGE compared to the LGE. On the other hand, in the *hs623KO* we observed a reduction of *Tcf12* expression that appeared to be specific to the SVZ and MZ of the MGE (Figure 3C and 3D). This result is consistent with the spatial activity of *hs623*, which is restricted to the SVZ of the MGE.

### Combined activity of POU and SOX TFs are required to maintain gene expression downstream of NKX2-1 in the MGE

To test the functional requirement of the LHX6 motifs in *hs623* we made site directed mutations that removed all four LHX6 motifs (hs623ΔLHX). In MGE primary cultures the activity of hs623ΔLHX was reduced by half when compared to the non-mutated hs623 (hs623WT, Figure 4A and 4B). Together, these experiments provide evidence that *hs623* activity, in part, depends on LHX6 and LHX8 and that there are additional TFs and signaling pathways required for the activity of *hs623*. Our earlier motif analysis of aREs discovered an enrichment of additional motifs such as HD-binding motifs (POU6f2 and HOX), SOX motifs and E-boxes (Figure 1E). To identify additional TF pathways responsible for the activity of *hs623* we looked at the other identified *de novo* motifs within *hs623* (Figure 1D). Located in the center of *hs623* we found two octamers (bound by POU TFs), of which one is adjacent to a SOX motif. Octamers are known to pair with SOX motifs to form central functional units regulating development in various cell types [34–36]. Initially, we analyzed the activity of the two individual octamers by generating single mutations of the two motifs (Figure 4A, hs623ΔOCT1 and hs623ΔOCT2). Mutating octamer 1 (*hs623ΔOCT1*) caused a significant reduction of *hs623* activity in MGE primary cultures, whereas mutating octamer 2 (*hs623ΔOCT2*) had no significant effect on hs623 activity (Figure 4B). Octamer 2 is located 3bp from a SOX consensus motif (Figure 4A). To assess the requirement of this combined motif for *hs623* activity, we generated a compound mutant with a combined mutation of octamer 2 and the paired SOX motif (hs623 ΔOCT2 + SOX). Hs623ΔOCT2 + SOX showed a significantly reduced activity when compared to hs623WT as well as, the two individual single mutants, hs623 ΔOCT2 and hs623ΔSOX (Figure 4B).

**Figure 4.**
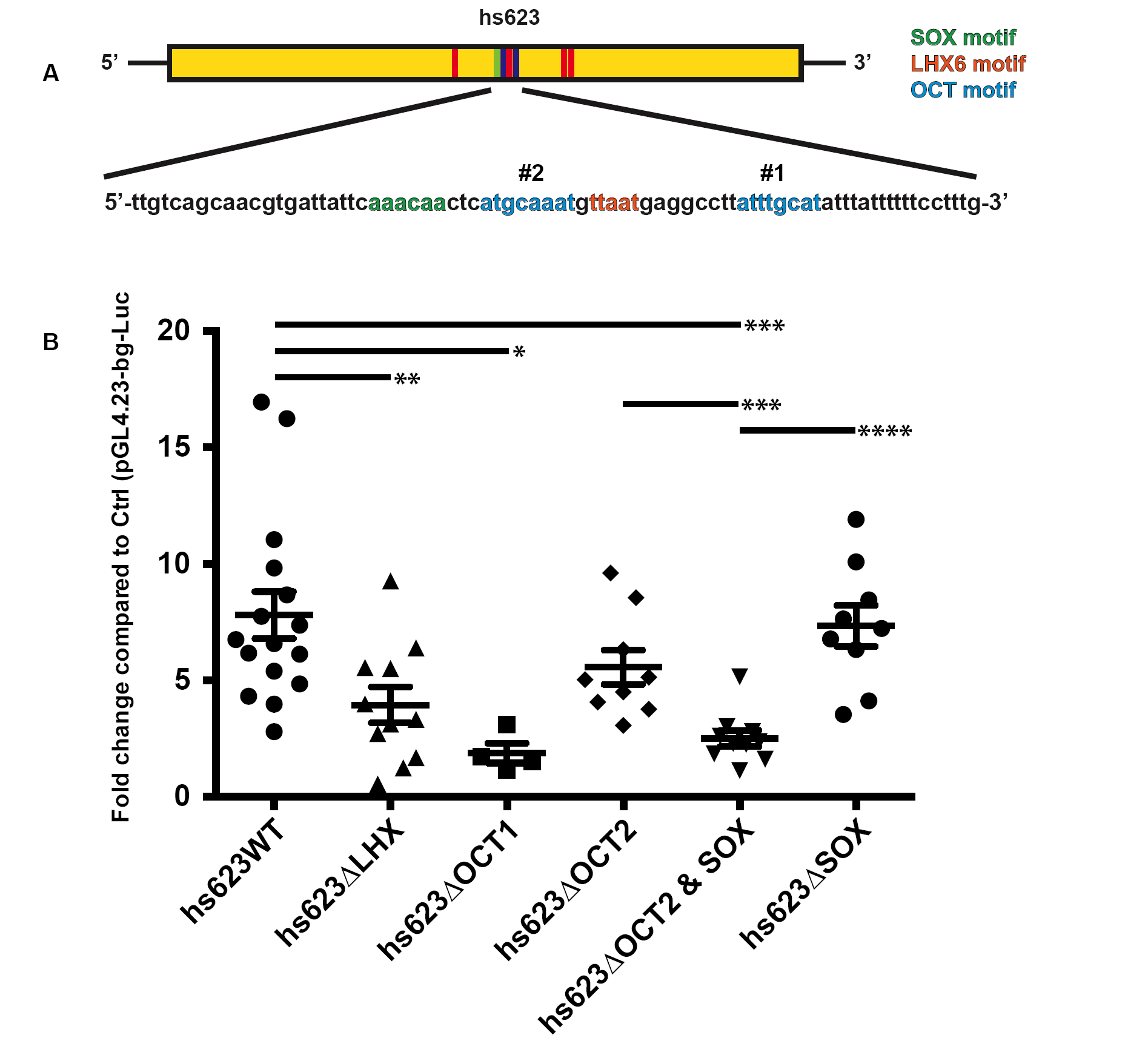
OCT and SOX motifs required for hs623 activity in primary MGE cultures. (A) Schematic of hs623 with LHX6, OCT and SOX motifs. (B) Luciferase reporter assay showing a reduced activity of hs623 when LHX6, OCT and SOX motifs in hs623 are mutated. Data are represented as mean ± SEM. Unpaired t-test was used to test significance between the groups: *p < 0.05, **p < 0.01, ***p < 0.001, ****p < 0.0001.

Altogether, our experiments show that *Tcf12* expression in the SVZ of the MGE is mediated, at least in part, through *hs623*, a RE that is strongly dependent on its OCT and SOX motifs and partially dependent on its LHX6 motifs. We have previously shown that gene expression in the SVZ of the MGE (including *Tcf12*) largely depends on NKX2-1 activity [18]. Existing mechanistic data show that NKX2-1 acts as a transcriptional repressor. Therefore, our findings suggest that the loss of *Tcf12* expression in the SVZ of the MGE *Nkx2-1c*KO is not due to the direct regulation of *Tcf12* by NKX2-1, but is a secondary effect due to the loss of additional TFs expressed genetically downstream of NKX2-1, including LHX6, LHX8, OCT and SOX TFs.

## DISCUSSION

Technical advancements in genome wide sequencing, chromosome capture and CRISPR/Cas9 technologies are increasing our understanding of genome organization. These data, combined with data showing RE activities *in vivo* (https://enhancer.lbl.gov/), TF binding and other epigenetic genomic data, and spatial gene expression data (http://www.brain-map.org/, http://www.eurexpress.org/ee/intro.html), are enabling the field to begin elucidating the genomic networks and the molecular mechanisms that direct brain development. Herein we have used many of these approaches to analyze gene expression in the developing mouse MGE. In the context of the *Nkx2-1*cKO mouse, our analysis of differential (WT vs. cKO) histone ChIP-seq data, and *de novo* sequence motif analysis, has provided evidence for additional TFs, REs, and signaling pathways that direct MGE development.

In this study, we showed that *Tcf12* expression in the SVZ of the MGE is mediated via hs623, an aRE bound by NKX2-1. The activity of hs623 and the expression of *Tcf12* depend on NKX2-1 activity, suggesting that NKX2-1 promotes *Tcf12* expression in this context. Surprisingly, we find no direct evidence showing that NKX2-1 activates *Tcf12* transcription via *hs623*. On the other hand, we show that LHX6, OCT and SOX motives are central to *hs623* activity. In fact, hs623 lacks NKX2-1 consensus motifs and its interaction with *hs623* can possibly be explained through binding to three flanking regions. Other alternative explanations are that NKX2-1 regulates *hs623* through either uncharacterized NKX2-1 motifs or through indirect binding to transcriptional complexes that bind *hs623*.

We have earlier demonstrated that NKX2-1 represses transcription in the MGE, similar to other NKX HD TFs that specify ventral parts of the developing neural tube [37,38]. Even at aREs, identified in the *Nkx2-1*cKO MGE, the NKX2-1 motifs mediate transcriptional repression, as exemplified by the intragenic *Tgfb3* RE in Sandberg et al. 2016 [18]. On the other hand, in the case of both the *Tgfb3* RE and *hs623*, LHX6 motifs promote enhancer activity. If NKX2-1 only represses transcription, it is unclear how loci such as LHX6 and LHX8 fail to be activated in the *Nkx2-1* mutants [16–18]. Furthermore it is unclear why NKX2-1 also binds loci that have reduced activity in the *Nkx2-1*cKO. These results suggest that, in some contexts, NKX2-1 may have an activating function. NKX2-1 binding to these loci might be required to keep them poised for subsequent activation by TFs and signaling pathways parallel and genetically downstream of NKX2-1, such as LHX, OCT, SOX and bHLH TFs. A similar model was presented in two studies looking at motor neuron development. In these cells, combinations of NEUROG2 (bHLH TF), LHX3, ISL-1, ONECUT1 and EBF direct the progression of the motor neuron fate through distinct sets of REs [8,9]. Similar to these models, we find an enrichment of LHX6 binding and e-boxes at aREs, a group of REs with a preferential activity in the SVZ of the MGE. This combination of TF binding and motif enrichment is not seen at NKX2-1 bound rREs, that have a relatively low MGE activity. These data highlight similarities in the molecular mechanisms that direct MGE and motor neuron development over time. In addition to combinatorial activity with other TFs, the activity of NKX2-1 might be affected by changes to chromatin modifications at specific loci over time.

The seemingly dual activity of NKX2-1 in the MGE is similar to its double-edged characteristics in regulating cancer development and progression. In this context, NKX2-1 has a role as lineage-survival oncogene in developing lung cancer tumors. On the other hand, NKX2-1 expression is also associated with a favorable prognosis in affected patients, due to its capacity to attenuate the invasive capacity of carcinomas [39]. Interestingly, this has been shown to be mediated through an abrogation of cellular response to TGFβ induced EMT, a signaling pathway that is directly repressed by NKX2-1 in the MGE [18,40]. By identifying the mechanisms through which NKX2-1 operate in the subpallial telencephalon we might also learn more about its enigmatic role in tumor biology. Our data provides evidence that LHX6 activates *Tcf12* expression through the *hs623* RE based on three observations: 1) LHX6 binding to *hs623 in vivo*, 2) a requirement of LHX6 motifs for hs623 activity in MGE cell culture assays, and 3) reduced expression of LHX6 in the *Nkx2-1*cKO.

In addition, our data show a combination of NKX2-1 and LHX6 binding to aREs that have an enrichment of E-box motifs. This combination of TFs (HD, LIM and E-box binding) is consistent with the transcriptional program governing the development and establishment of terminal effector gene expression in motor neurons [8,9]. These beautiful studies show that TFs are shuttled between different REs along the progression of motor neuron development. Our experimental setup lacks the temporal resolution to make these kinds of predictions. For the future, it would be interesting to know; 1) at what time point in the developing MGE (VZ, SVZ or MZ) are the various REs active and, 2) and the temporal pattern of TF binding at these REs. This would give us important information that could help us understand the activating and repressing mechanics through which NKX2-1, LHX6 and other TFs direct MGE development.

In this study we show an example of an aRE (hs623) that required for maintaining normal levels of *Tcf12* expression in the SVZ and MZ of the MGE *in vivo*. The activity of the RE hs623 depends on two octamers, providing evidence that OCT TFs are central to MGE development.

OCT TFs are important regulators of stem cell maintenance and the progression of neurogenesis. OCT4 is central for propagating undifferentiated embryonic stem cells and has the ability to induce pluripotent stem cells from embryonic and adult fibroblasts [41,42]. On the contrary, BRN2, together with Ascl1 and Myt1l, can efficiently trans-differentiate embryonic and postnatal fibroblasts into functional neurons [43]. BRN1, BRN2 and OCT6 mutants show defects in layering of the neocortex, due to their role in initiating radial migration of cortical projection neurons, further highlighting their role in promoting neurogenesis [44–46]. Furthermore, we find an enrichment of both octamers and E-boxes in REs promoting gene expression in the MGE, suggesting that the TF machinery directing trans-differentiation of fibroblasts into neurons is similar to the TF machinery inducing neuronal phenotypes in the MGE. Trans-differentiating fibroblast to neurons using Brn2, Ascl1 and Myt1l generate cells with a mixed neuronal phenotype [43], indicating that these TFs are required for promoting the neuronal fate, without any preference for specific neuronal lineages. Taken together, this suggests a model where neuronal fate and phenotype is directed through separate, although integrated, TF pathways in the MGE.

One of the octamers in hs623 is paired with a SOX motif. The SOX TF family consists of a large number of genes that direct embryonic development and cell differentiation. They bind loosely to the minor groove of the DNA and their target gene specificity is guided through the interaction with cell type specific partner factors such as OCT TFs. The combined activity of SOX2 and different OCT TFs are important regulators of gene expression in undifferentiated embryonic stem cells (ESCs) and neural progenitor cells (NPCs) [47]. SOX2 and OCT4 (POU5F1) bind REs in ECSs, whereas SOX2 and BRN2 (POU3F2) co-occupy REs in NPCs [35,47–49]. SOX and OCT motifs have also been shown to direct transcription in both ESCs and NPCs in the forebrain [35,36]. Today we do not know what specific OCT and SOX TFs that are required to activate transcription in the SVZ of the MGE, via REs like hs623, BRN1 (POU3F3), BRN2 (POU3F2), BRN4 (POU3F4), BRN5 (POU6F1) and OCT6 (POU3F1) are all expressed here, but little is know regarding their function in this context. A large number of different SOX TFs are expressed in the MGE and several of them show a reduced expression in the *Nkx2-1*cKO, such as Sox1, Sox2, Sox6, Sox11 and Sox21 constituting possible candidates for promoting *Tcf12* expression via hs623. Sox6 is required for patterning of the subpallium and generation on MGE derived interneurons [50,51], but when looking at *Tcf12* expression in the E13.5 MGE of a conditional *Sox6* mouse [52] with an *Nkx2-1*-Cre line [53], we found no significant change in *Tcf12* expression (Additional file 2). From this we can exclude that SOX6 is not sufficient for promoting *Tcf12* expression. Further studies should be performed to identify the specific OCT and SOX TFs directing transcriptional activation in the MGE.

Here, in our new analysis of the *Nkx2-1c*KO, we found a large number aREs. Some of these are near the loci of the *Tcf4* and *Tcf12* bHLH TF encoding genes. The *Nkx2-1*cKO shows a near complete loss of *Tcf12* expression in the SVZ and MZ of the MGE. We found an aRE intronic to *Tcf12* (*hs623*) that has activity in the SVZ and MZ of the MGE (Figure 2E). Deletion of *hs623* leads to a reduced *Tcf12* expression in the VZ and MZ of the MGE. This result suggests that *Tcf12* expression is regulated through several aREs, including *hs623*, and that there is redundancy between these REs. Enhancer redundancy has been demonstrated in the developing telencephalon and limb where REs sharing a similar spatiotemporal activity provides robustness to gene expression [54,55]. We also find that there are different genetic programs directing *Tcf12* expression in various cell types of the MGE. *Tcf12* expression is initiated in the VZ of the MGE; this expression is largely unaffected in the *Nkx2-1*cKO, indicating that *Tcf12* expression in this region is not mediated through hs623 and largely NKX2-1 independent.

Altogether, these data provide evidence of transcriptional circuitry that connects the initiation of MGE fate in the VZ by *Nkx2-1* and *Otx2*, to the maturation of cells in the SVZ and MZ by driven through REs such as *hs632,* whose activity integrates signals from LHX, OCT, SOX and bHLH TFs [16,18,56]. Future studies will investigate how TFs, chromatin-binding,-reading and-remodeling proteins integrate to direct GABAergic and cholinergic development in the subpallial telencephalon.

## CONCLUSION

In our study we use a combination of genomics, CRISPR / Cas9 engineering and TF motif analysis to investigate the transcriptional networks guiding development of the MGE and its descendants. Whereas NKX2-1 is required for initiating MGE characteristics in the VZ, we provide evidence that a combination of LHX, OCT, SOX and bHLH TFs are central for maintaining gene expression in the SVZ and MZ, genetically down-stream of NKX2-1. Here we generate a mouse mutant in whom we delete a *Tcf12* intragenic RE, showing its requirement for maintaining *Tcf12* transcription in the SVZ and MZ of the MGE. The activity of this *Tcf12* enhancer, in primary cultures of MGE cells, largely depends on an octamer and a combined octamer and SOX motif. Altogether, our study identifies a genomic framework through which a combination of LHX, OCT, SOX and bHLH TFs direct MGE differentiation through the expression terminal effector genes.

## LIST OF ABBREVIATIONS

TF: transcription factor; REs: regulatory elements; aREs: activating regulatory element; E: embryonic day; MGE: medial ganglionic eminence; rREs: repressing regulatory element; bHLH: basic-helix-loop-helix; MZ: mantle zone; SVZ: sub-ventricular zone; HD: homeodomain; GO: gene ontology; LGE: lateral ganglionic eminence; VZ: ventricular zone; WT: wild-type; cKO: conditional knockout

## DECLARATIONS

### Ethics approval and consent to participate

All animal procedures were performed following protocols (AN143392-02B [J.L.R.R.] and AN171299-01 [B.L.B.]) authorized by the University of California San Francisco Institutional Animal Care and Use Committee (IACUC).

### Consent for publication

Not applicable.

### Availability of data and material

The datasets supporting the conclusions of this article are available in the NCBI’s GEO repository, GEO Series accession number GSE85705 (https://www.ncbi.nlm.nih.gov/geo/query/acc.cgi?acc=GSE85705). Additional material is available from the corresponding author upon request.

### Authors’ contributions

M.S., J.L.R.R. and A.N. conceived the experiments, interpreted the results, wrote, reviewed and edited the manuscript. M.S. designed and performed all experiments. J.H. and B.B. injected sgRNA and Cas9 to generate *hs623*KO mice. L.T. performed the differential ChIP-seq enrichment analysis.

## Acknowledgements

We thank members of the Rubenstein, Taher, Black and Nord laboratories for advice and critical evaluation of the data during the course of the study.

## FIGURE LEGENDS

Additional file 1. Identified activating (sheet “aRE”) and repressing (sheet “rRE”) regulatory elements in Nkx2-1cKO MGE at e13.5. *In vivo* activity of aREs (sheet “VISTA aRE”) and rREs (sheet “ VISTA rRE”) VISTA transgenics at E11.5.

Additional file 2. In situ analysis of *Tcf12* in WT (A) and *Sox6* conditional knockout (B) forebrain at E13.5.

## REFERENCES

1. Nord AS, Pattabiraman K, Visel A, Rubenstein JLR. Genomic Perspectives of Transcriptional Regulation in Forebrain Development. Neuron. Elsevier Inc; 2015;85:27–47.

2. De Rubeis S, He X, Goldberg AP, Poultney CS, Samocha K, Ercument Cicek A, et al. Synaptic, transcriptional and chromatin genes disrupted in autism. 2014.

3. Pennacchio LA, Ahituv N, Moses AM, Prabhakar S, Nobrega MA, Shoukry M, et al. In vivo enhancer analysis of human conserved non-coding sequences. Nature. Nature Publishing Group; 2006;444:499–502.

4. Visel A, Blow MJ, Li Z, Zhang T, Akiyama JA, Holt A, et al. ChIP-seq accurately predicts tissue-specific activity of enhancers. Nature Publishing Group; 2009;457:854–8.

5. Zinzen RP, Girardot C, Gagneur J, Braun M, Furlong EEM. Combinatorial binding predicts spatio-temporal cis-regulatory activity. Nature Publishing Group; 2009;462:65–70.

6. Bonev B, Cohen NM, Szabo Q, Fritsch L, Papadopoulos GL, Lubling Y, et al. Multiscale 3D Genome Rewiring during Mouse Neural Development. Cell. Elsevier; 2017;171:557–572.e24.

7. Visel A, Taher L, Girgis H, May D, Golonzhka O, Hoch RV, et al. A high-resolution enhancer atlas of the developing telencephalon. Cell. 2013;152:895–908.

8. Rhee HS, Closser M, Guo Y, Bashkirova EV, Tan GC, Gifford DK, et al. Expression of Terminal Effector Genes in Mammalian Neurons Is Maintained by a Dynamic Relay of Transient Enhancers. Neuron. Elsevier; 2016;92:1252–65.

9. Velasco S, Ibrahim MM, Kakumanu A, Garipler G, Aydin B, Al-Sayegh MA, et al. A Multi-step Transcriptional and Chromatin State Cascade Underlies Motor Neuron Programming from Embryonic Stem Cells. Cell Stem Cell. 2016.

10. Lupiáñez DG, Kraft K, Heinrich V, Krawitz P, Brancati F, Klopocki E, et al. Disruptions of topological chromatin domains cause pathogenic rewiring of gene-enhancer interactions. Cell. 2015;161:1012–25.

11. Campbell K. Dorsal-ventral patterning in the mammalian telencephalon. Curr. Opin. Neurobiol. 2003;13:50–6.

12. Rubenstein JLR, Campbell K. Neurogenesis in the Basal Ganglia. Patterning and Cell Type Specification in the Developing CNS and PNS. Elsevier; 2013. pp. 455–73.

13. Zhao Y, Marin O, Hermesz E, Powell A, Flames N, Palkovits M, et al. The LIM-homeobox gene Lhx8 is required for the development of many cholinergic neurons in the mouse forebrain. Proc. Natl. Acad. Sci. U.S.A. 2011;100:9005–10.

14. Fragkouli A, van Wijk NV, Lopes R, Kessaris N, Pachnis V. LIM homeodomain transcription factor-dependent specification of bipotential MGE progenitors into cholinergic and GABAergic striatal interneurons. Development. 2009;136:3841–51.

15. Du T, Xu Q, Ocbina PJ, Anderson SA. NKX2.1 specifies cortical interneuron fate by activating Lhx6. Development [Internet]. 2008;135:1559–67. Available from: http://eutils.ncbi.nlm.nih.gov/entrez/eutils/elink.fcgi?dbfrom=pubmed&id=1833967 4&retmode=ref&cmd=prlinks

16. Sussel L, Marin O, Kimura S, Rubenstein JL. Loss of Nkx2.1 homeobox gene function results in a ventral to dorsal molecular respecification within the basal telencephalon: evidence for a transformation of the pallidum into the striatum. Development. 1999.

17. Butt SJB, Sousa VH, Fuccillo MV, Hjerling-Leffler J, Miyoshi G, Kimura S, et al. The Requirement of Nkx2-1 in the Temporal Specification of Cortical Interneuron Subtypes. Neuron. 2008;59:722–32.

18. Sandberg M, Flandin P, Silberberg S, Su-Feher L, Price JD, Hu JS, et al. Transcriptional Networks Controlled by NKX2-1 in the Development of Forebrain GABAergic Neurons. Neuron. 2016;91:1260–75.

19. Vogt D, Hunt RF, Mandal S, Sandberg M, Silberberg SN, Nagasawa T, et al. Lhx6 directly regulates Arx and CXCR7 to determine cortical interneuron fate and laminar position. Neuron. 2014;82:350–64.

20. Kusakabe T, Kawaguchi A, Hoshi N, Kawaguchi R, Hoshi S, Kimura S. Thyroid-specific enhancer-binding protein/NKX2.1 is required for the maintenance of ordered architecture and function of the differentiated thyroid. Mol. Endocrinol. 2006;20:1796–809.

21. Schüller U, Heine VM, Mao J, Kho AT, Dillon AK, Han Y-G, et al. Acquisition of Granule Neuron Precursor Identity Is a Critical Determinant of Progenitor Cell Competence to Form Shh-Induced Medulloblastoma. Cancer Cell. 2008;14:123–34.

22. Madisen L, Zwingman TA, Sunkin SM, Oh SW, Zariwala HA, Gu H, et al. A robust and high-throughput Cre reporting and characterization system for the whole mouse brain. Nat Neurosci. Nature Publishing Group; 2010;13:133–40.

23. Wang H, Yang H, Shivalila CS, Dawlaty MM, Cheng AW, Zhang F, et al. One-Step Generation of Mice Carrying Mutations in Multiple Genes by CRISPR/Cas-Mediated Genome Engineering. Cell. 2013;153:910–8.

24. De Val S, Anderson JP, Heidt AB, Khiem D, Xu S-M, Black BL. Mef2c is activated directly by Ets transcription factors through an evolutionarily conserved endothelial cell-specific enhancer. Dev. Biol. 2004;275:424–34.

25. Zhao Y, Flandin P, Long JE, Cuesta MD, Westphal H, Rubenstein JLR. Distinct molecular pathways for development of telencephalic interneuron subtypes revealed through analysis of Lhx6 mutants. J. Comp. Neurol. 4. 2008;510:79–99.

26. Schaeren-Wiemers N, Gerfin-Moser A. A single protocol to detect transcripts of various types and expression levels in neural tissue and cultured cells: in situ hybridization using digoxigenin-labelled cRNA probes. Histochemistry. Springer-Verlag; 1993;100:431–40.

27. Schmittgen TD, Livak KJ. Analyzing real-time PCR data by the comparative CT method. Nat Protoc. 2008;3:1101–8.

28. Livak KJ, Schmittgen TD. Analysis of relative gene expression data using real-time quantitative PCR and the 2(-Delta Delta C(T)) Method. Methods. Academic Press; 2001;25:402–8.

29. Thomas-Chollier M, Darbo E, Herrmann C, Defrance M, Thieffry D, van Helden J. A complete workflow for the analysis of full-size ChIP-seq (and similar) data sets using peak-motifs. Nat Protoc. 2012;7:1551–68.

30. Silberberg SN, Taher L, Lindtner S, Sandberg M, Nord AS, Vogt D, et al. Subpallial Enhancer Transgenic Lines: a Data and Tool Resource to Study Transcriptional Regulation of GABAergic Cell Fate. Neuron. Elsevier; 2016;92:59–74.

31. Hu JS, Vogt D, Sandberg M, Rubenstein JL. Cortical interneuron development: a tale of time and space. Development. Oxford University Press for The Company of Biologists Limited; 2017;144:3867–78.

32. McLean CY, Bristor D, Hiller M, Clarke SL, Schaar BT, Lowe CB, et al. GREAT improves functional interpretation of cis-regulatory regions. Nat Biotechnol. 2010;28:495–501.

33. Visel A, Minovitsky S, Dubchak I, Pennacchio LA. VISTA Enhancer Browser--a database of tissue-specific human enhancers. Nucleic Acids Res. 2007;35:D88–92.

34. Kamachi Y, Uchikawa M, Kondoh H. Pairing SOX off: with partners in the regulation of embryonic development. Trends in Genetics. 2000;16:182–7.

35. Josephson R, Muller T, Pickel J, Okabe S, Reynolds K, Turner PA, et al. POU transcription factors control expression of CNS stem cell-specific genes. Development. 1998;125:3087–100.

36. Bery A, Martynoga B, Guillemot F, Joly J-S, Rétaux S. Characterization of Enhancers Active in the Mouse Embryonic Cerebral Cortex Suggests Sox/Pou cis-Regulatory Logics and Heterogeneity of Cortical Progenitors. Cereb. Cortex. Oxford University Press; 2014;24:2822–34.

37. Briscoe J, Pierani A, Jessell TM, Ericson J. A homeodomain protein code specifies progenitor cell identity and neuronal fate in the ventral neural tube. Cell. 2000;101:435–45.

38. Muhr J, Andersson E, Persson M, Jessell TM, Ericson J. Groucho-mediated transcriptional repression establishes progenitor cell pattern and neuronal fate in the ventral neural tube. Cell. 2001.

39. Yamaguchi T, Hosono Y, Yanagisawa K, Takahashi T. NKX2-1/TTF-1: An Enigmatic Oncogene that Functions as a Double-Edged Sword for Cancer Cell Survival and Progression. Cancer Cell. Elsevier; 2013;23:718–23.

40. Saito R-A, Watabe T, Horiguchi K, Kohyama T, Saitoh M, Nagase T, et al. Thyroid Transcription Factor-1 Inhibits Transforming Growth Factor-β–Mediated Epithelial-to-Mesenchymal Transition in Lung Adenocarcinoma Cells. Cancer Res. American Association for Cancer Research; 2009;69:2783–91.

41. Boyer LA, Lee TI, Cole MF, Johnstone SE, Levine SS. Core transcriptional regulatory circuitry in human embryonic stem cells. Cell. 2005.

42. Takahashi K, Yamanaka S. Induction of pluripotent stem cells from mouse embryonic and adult fibroblast cultures by defined factors. Cell. 2006;126:663–76.

43. Vierbuchen T, Ostermeier A, Pang ZP, Kokubu Y, Südhof TC, Wernig M. Direct conversion of fibroblasts to functional neurons by defined factors. Nature. 2010;463:1035–41.

44. McEvilly RJ, de Diaz MO, Schonemann MD, Hooshmand F, Rosenfeld MG. Transcriptional regulation of cortical neuron migration by POU domain factors. Science. 2002;295:1528–32.

45. Sugitani Y, Nakai S, Minowa O, Nishi M, Jishage K-I, Kawano H, et al. Brn-1 and Brn-2 share crucial roles in the production and positioning of mouse neocortical neurons. Genes & Development. Cold Spring Harbor Lab; 2002;16:1760–5.

46. Dominguez MH, Ayoub AE, Rakic P. POU-III Transcription Factors (Brn1, Brn2, and Oct6) Influence Neurogenesis, Molecular Identity, and Migratory Destination of Upper-Layer Cells of the Cerebral Cortex. Cereb. Cortex. Oxford University Press; 2013;23:2632–43.

47. Lodato MA, Ng CW, Wamstad JA, Cheng AW, Thai KK, Fraenkel E, et al. SOX2 co-occupies distal enhancer elements with distinct POU factors in ESCs and NPCs to specify cell state. Barsh GS, editor. PLoS Genet. Public Library of Science; 2013;9:e1003288.

48. Ambrosetti DC, Basilico C, Dailey L. Synergistic activation of the fibroblast growth factor 4 enhancer by Sox2 and Oct-3 depends on protein-protein interactions facilitated by a specific spatial arrangement of factor binding sites. Molecular and Cellular Biology. American Society for Microbiology; 1997;17:6321–9.

49. Nishimoto M, Fukushima A, Okuda A, Muramatsu M. The gene for the embryonic stem cell coactivator UTF1 carries a regulatory element which selectively interacts with a complex composed of Oct-3/4 and Sox-2. Molecular and Cellular Biology. American Society for Microbiology (ASM); 1999;19:5453–65.

50. Azim E, Jabaudon D, Fame RM, Macklis JD. SOX6 controls dorsal progenitor identity and interneuron diversity during neocortical development. Nat Neurosci. NIH Public Access; 2009;12:1238–47.

51. Batista-Brito R, Rossignol E, Hjerling-Leffler J, Denaxa M, Wegner M, Lefebvre V, et al. The cell-intrinsic requirement of Sox6 for cortical interneuron development. Neuron. 2009;63:466–81.

52. Dumitriu B, Dy P, Smits P, Lefebvre V. Generation of mice harboring aSox6 conditional null allele. genesis. 2006;44:219–24.

53. Xu Q, Tam M, Anderson SA. Fate mapping Nkx2.1-lineage cells in the mouse telencephalon. J. Comp. Neurol. 2008;506:16–29.

54. Osterwalder M, Barozzi I, Tissières V, Fukuda-Yuzawa Y, Mannion BJ, Afzal SY, et al. Enhancer redundancy provides phenotypic robustness in mammalian development. Nature. Nature Publishing Group; 2018;489:57.

55. Dickel DE, Ypsilanti AR, Pla R, Zhu Y, Barozzi I, Mannion BJ, et al. Ultraconserved Enhancers Are Required for Normal Development. Cell. 2018;172:491–499.e15.

56. Hoch RV, Lindtner S, Price JD, Rubenstein JLR. OTX2 Transcription Factor Controls Regional Patterning within the Medial Ganglionic Eminence and Regional Identity of the Septum. Cell Rep. 2015;12:482–94.

